# Ceramide Synthases Regulate Myristate-Induced Intestinal IRE1α Activation

**DOI:** 10.64898/2026.05.28.728542

**Authors:** Chelsea L Doll, Mary R Gordon, Marco Padilla-Rodriguez, Eugene Jap, Paul A Boasiako, Marilyn T Marron, Brandon K Dahl, Keila S Espinoza, Drew M Seiser, Rachel J Ren, Curtis A Thorne, Justin M Snider, Ashley J Snider

## Abstract

**Background & Aims:** High-fat diets (HFDs) are a major modifiable risk factor for intestinal health. Current research focuses primarily on palmitate (C16:0); however, myristate (C14:0, rich in dairy products) has been minimally investigated. HFDs increase ceramide generation which drives endoplasmic reticulum (ER) stress; with both sphingolipids and ER stress being key contributors to intestinal biology. Whether different fatty acids uniquely impact sphingolipid metabolism and ER stress in intestinal biology has not been well defined.

**Methods:** Human colon epithelial cells were utilized to determine the role of ceramide synthases (CerS) 5 and 6 on myristate-induced ER stress using pharmacologic inhibitors and siRNA. Intestinal epithelial cell specific CerS5 and/or CerS6 knockout mice of both sexes were fed a control, high milk-fat, or high lard-fat diet for 16 weeks. Cells and colon tissues were analyzed for lipids, mRNA, and protein.

**Results:** Myristate treatment increased C14:0-ceramide and induced IRE1α-dependent ER stress. Inhibition of CerS suppressed these effects, yet knockdown of CerS5/6, the primary enzymes generating C14:0-ceramide, unexpectedly exacerbated IRE1α activation both *in vitro* and *in vivo*, potentially due to depletion of dihydro(dh)sphingosine.

**Conclusions:** CerS are required for myristate-induced IRE1α activation and restoration of the sphingoid base pool provides partial protection from intestinal ER stress.

**SYNOPSIS:** This study identifies a new mechanism linking dietary fats to intestinal cell stress. Ceramide synthases drive ER stress triggered by myristate, a dairy-derived fat, while restoring sphingoid bases partially protects cells, revealing a new role for sphingolipids in shaping intestinal responses to diet.

**Graphical abstract:** 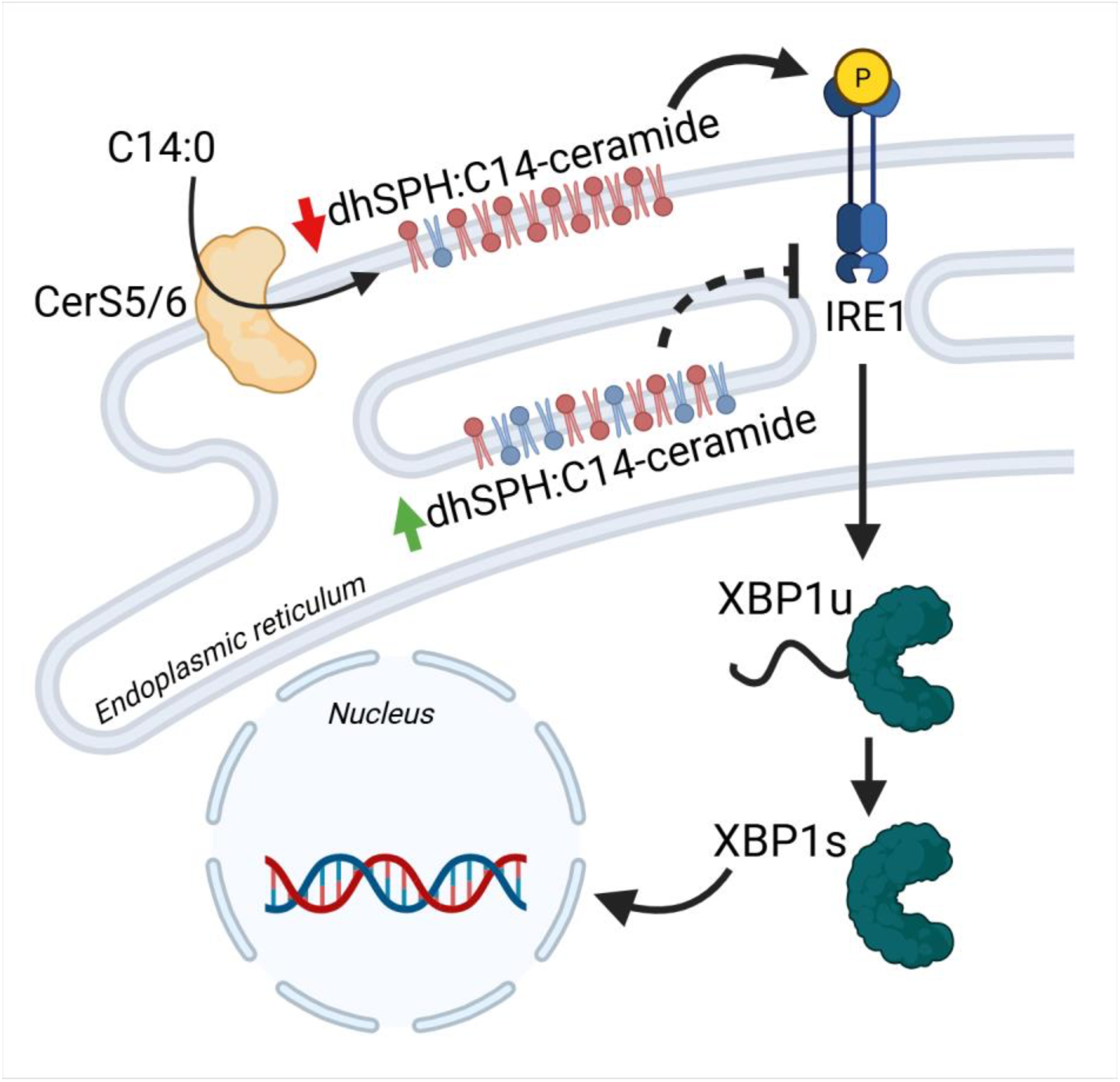

## BACKGROUND

Rates of inflammatory bowel disease (IBD) are increasing at an alarming rate worldwide [1–5]. Quality of life is severely diminished in patients with IBD due to symptoms of chronic abdominal pain, diarrhea, bloody stool, weight loss, and fatigue. Further, those afflicted with IBD are at an increased risk of developing early-onset colorectal cancer [6]. Although causes of IBD are multifactorial, diet is one of the major modifiable risk factors and a potential avenue for nutritional and therapeutic intervention [7–10]. Understanding how diet impacts cellular biology in the intestines is crucial in developing effective avenues for prevention of IBD, as well as diagnostic markers and improved treatment options.

Sphingolipids are bioactive lipids involved in many fundamental processes including differentiation, senescence, apoptosis, and cell cycle-arrest [11], and have been implicated as regulators in intestinal diseases such as IBD, colon cancer, and colitis-associated colon cancer [12–18]. Ceramides are of particular interest as they are central to multiple sphingolipid metabolic pathways. Both exogenous and endogenous fatty acids are incorporated into ceramides using two key enzymes: serine palmitoyltransferase (SPT) and ceramide synthase (CerS). SPT is the first step in *de novo* sphingolipid generation and canonically condenses serine and palmitate, a 16-carbon saturated fatty acid, to create an 18-carbon sphingoid base. CerS, of which there are six isoforms, add an additional fatty acid into the acyl-chain to generate ceramide. Each CerS exhibits a preference for a fatty acyl chain length [19]. Of interest, CerS5 and CerS6 preferentially utilize C14:0 and C16:0 fatty-acyl CoAs yielding C14:0 and C16:0 ceramide. In metabolic disease, these differences in chain length are appreciated for having distinct effects on cellular outcomes [20–22]; however, whether this extends to intestinal tissues has not been fully investigated.

Although high fat diets (HFDs) are well characterized for promotion of systemic inflammation [23], in the intestines this effect is compounded due to altered lipid metabolism, as well as increased ER stress and gut permeability [24–27]. Investigation into the effect of dietary lipids on human physiology has, until recently, focused on saturated versus unsaturated fatty acids, with saturated fats being associated with poorer health outcomes. This duality has also been mirrored in sphingolipid metabolism whereby excessive intake of saturated fatty acids, not unsaturated, in healthy adults for eight-weeks increased serum ceramide levels [28]. Dietary fatty acids; however, vary in chain length depending on the foods consumed. Most of the literature to date has primarily focused on palmitate (C16:0), which is found in red meat and processed foods. Milk-based products, on the other hand, have high amounts of the saturated fatty acid myristate (C14:0) which has been minimally investigated. Intake of myristate has been associated with increased risk of IBD relapse [29], while other studies have inversely associated dairy consumption with IBD diagnosis [30 31]. The nebulous role of dairy, and more specifically the fatty acid myristate, in IBD led us to examine its impact on intestinal biology.

Endoplasmic reticulum (ER) stress is well established as a risk factor for IBD [32–36]. There are three evolutionary conserved pathways activated during ER stress (IRE1α, PERK, and ATF6) that target degradation of mis-folded proteins, halt generation of new proteins and lipids, and if left unresolved lead to inflammation and apoptosis. In addition to recognizing proteotoxic stress, these pathways are also responsive to perturbations in membrane lipids [37 38]. Excess saturated fatty acids and a subsequent increase in sphingolipid generation, particularly ceramide, has been shown to induce ER stress in multiple tissues [39–42]. Although ER stress and sphingolipids have been implicated in IBD, the interplay of the two in intestinal biology is not well understood.

Our previously published work in rat small intestinal epithelial cells (IEC6) and intestines from C57BL/6 mice fed a milk-fat based diet (MFBD) demonstrated that myristate increases intestinal ER stress [13]. Here we report that CerS are required for myristate-induced IRE1α activation and that restoration of the sphingoid base pool may provide protection from intestinal ER stress.

## RESULTS

### Myristate induces IRE1α-dependent ER stress

Our previous studies in IEC6 cells demonstrated that myristate activated ER stress in a CerS5/6 dependent manner. To investigate the role of myristate on ER stress in the colon we treated human colon epithelial cells (HCEC 1CT) with vehicle or increasing doses of myristate for up to 24 h as previously described [43–45]. Myristate treatment increased expression and phosphorylation of IRE1α in a dose dependent manner; however, phospho-EIF and grp78 expression, downstream of PERK and ATF6 respectively, did not change (Figure 1A). To assess signals downstream of IRE1α we examined XBP1 splicing (XBP1s) after myristate treatment, using tunicamycin as a positive control. XBP1s increased significantly following 16 h of myristate treatment with a maximal response at 24 h for both myristate and tunicamycin (Figure 1B, Supplemental Figure 1A). The mRNA expression of downstream ER stress markers, ERDJ4 and CHOP, were also significantly elevated starting at 16 h (Figure 1C, Supplemental Figure 1B). In addition, IL-6, a cytokine correlated with IBD, and previously shown to be downstream of IRE1α-XBP1 increased starting at 8 h (Figure 1C). Cell viability was not affected by myristate treatment (Supplemental Figure 1C). Myristate treatment in a second human colon epithelial cell line (HCEC 2CT) also increased XBP1s, as well as mRNA expression of ERDJ4, CHOP, and IL-6 (Supplemental Figure 1D, 1E). To further elucidate the role of IRE1α in myristate-induced ER stress we used an inhibitor of IRE1α (4µ8C) and generated HCEC^ΔIRE1α^ cell lines using CRISPR-Cas9. The addition of 4µ8C abrogated myristate-induced XBP1s and upregulation of ERDJ4, but not CHOP, expression (Figure 1D, 1E). Similarly, loss of IRE1α decreased both tunicamycin and myristate-induced XBP1s and ERDJ4 (Figure 1F, 1G). Loss of IRE1α increased CHOP expression after myristate treatment (Figure 1G), suggesting that CHOP expression is not downstream of IRE1α in response to myristate, and that loss of IRE1α may lead to compensation of other ER stress pathways in response to myristate. Collectively, these data demonstrate that myristate induces IRE1α-dependent ER stress in a dose and time-dependent manner.

**Figure 1:**
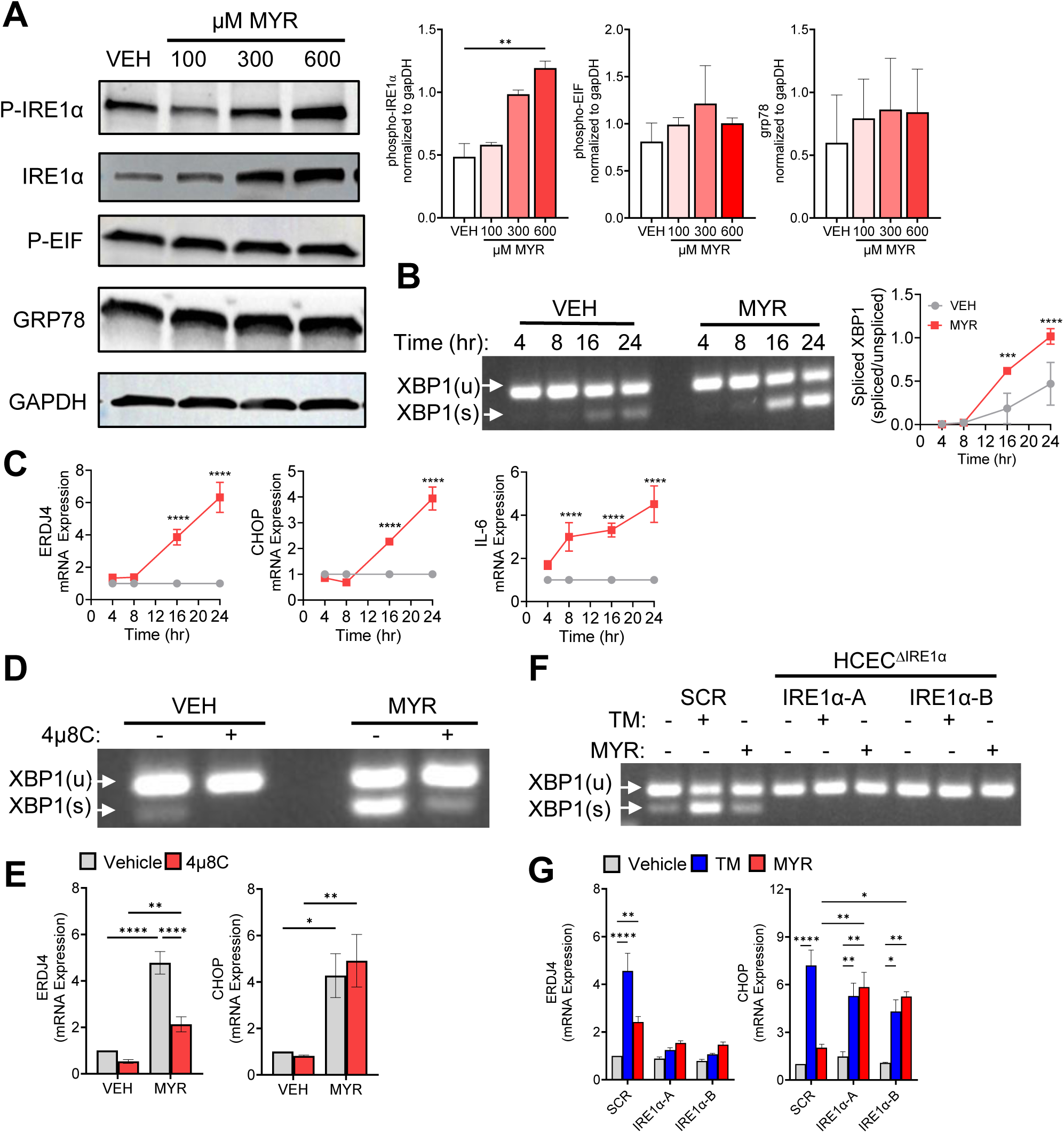
Myristate induces IRE1α-dependent ER stress. **(A)** HCEC-1CT cells were treated with vehicle (VEH, 2% FAF BSA) or indicated doses of myristate (MYR) for 24 h. Western blot of phosphorylated-IREα, total IREα, phosphorylated-EIF, GRP78, and GAPDH as a loading control with quantification (right). **(B&C)** HCEC-1CT were treated with 600µM MYR or VEH for indicated times. **(B)** XBP1 splicing; using extracted RNA and PCR primers to amplify unspliced (XBP1u) and spliced (XBP1s) with quantification (right). **(C)** mRNA levels of ERJD4, CHOP, and IL-6 analyzed by RT-qPCR and normalized to RPLPO. **(D&E)** HCEC 1CT cells were treated with 600µM MYR or VEH with or without 100µM IRE1α inhibitor 4µ8C for 24 h. **(D)** XBP1 splicing and **(E)** mRNA expression of ERDJ4 and CHOP. **(F&G)** IREα knockout using CRISPR/Cas9 in HCEC 1CT. SCR – scramble. IREα-A&B – monoclonal IREα CRISPR cell lines. Cells were treated with 600µM MYR, tunicamycin (TM), or VEH for 24 h. **(F)** XBP1 splicing and **(G)** mRNA expression of ERDJ4 and CHOP. Data represent mean ± SEM, n=3, *p<0.05, **p<0.01, ***p<0.001, ****p<0.0001.

### Myristate increases C14:0 ceramide in the ER

HFDs and/or saturated fatty acid treatment have been shown to induce ER stress in multiple tissues through increased generation of sphingolipids, and our previous work demonstrated that myristate increased generation of C14:0 ceramide. Therefore, we treated HCEC 1CT with an isotope labeled myristate and measured incorporation into sphingolipids using LC-MS/MS (Figure 2A). Treatment with labeled myristate resulted in a significant increase in labeled C14:0 ceramide and C14:0 hexosylceramide (ISO C14:0) (Figure 2B). Labeled myristate was minimally incorporated into C14:0 dihydro(dh)ceramide and sphingomyelin (Figure 2B). Comparison of labeled myristate into sphingolipids demonstrated that C14:0 ceramide contained the most C14:0 label as compared to dhceramide, hexosylceramide, and sphingomyelin at 24 h (Figure 2C). Although we measured increased incorporation of myristate into C14:0 sphingolipids, levels of total dhceramide, ceramide, and hexosylceramide levels remained unchanged over the time course (Figure 2D), whereas total sphingomyelin levels decreased after myristate treatment (Figure 2D). Interestingly, *de novo* sphingolipid generation increased over time in vehicle treated cells, as measured through increased d18:0 dhsphingosine (Figure 2E); however, myristate-treatment halted *de novo* sphingolipid synthesis (Figure 2E) leading to a decrease in the ratio of d18:0 dhsphingosine to C14:0 ceramide (Figure 2F). To elucidate if myristate and ceramide are localizing and accumulating in the ER we performed click chemistry by treating HCEC 1CT with 600µM of alkyne-modified myristate and imaged via confocal microscopy. Colocalization of ceramide with myristate (Figure 2G) and calnexin (Figure 2H) was increased after 16 h compared to vehicle treated cells, though the colocalization of myristate and calnexin did not change over time (data not shown). This increased colocalization at 16 h coincided with the myristate-induced ER stress previously described (Figure 1B, 1C). These data suggest that myristate decreases *de novo* sphingolipid synthesis and increases levels and localization of C14:0 ceramide in the ER.

**Figure 2:**
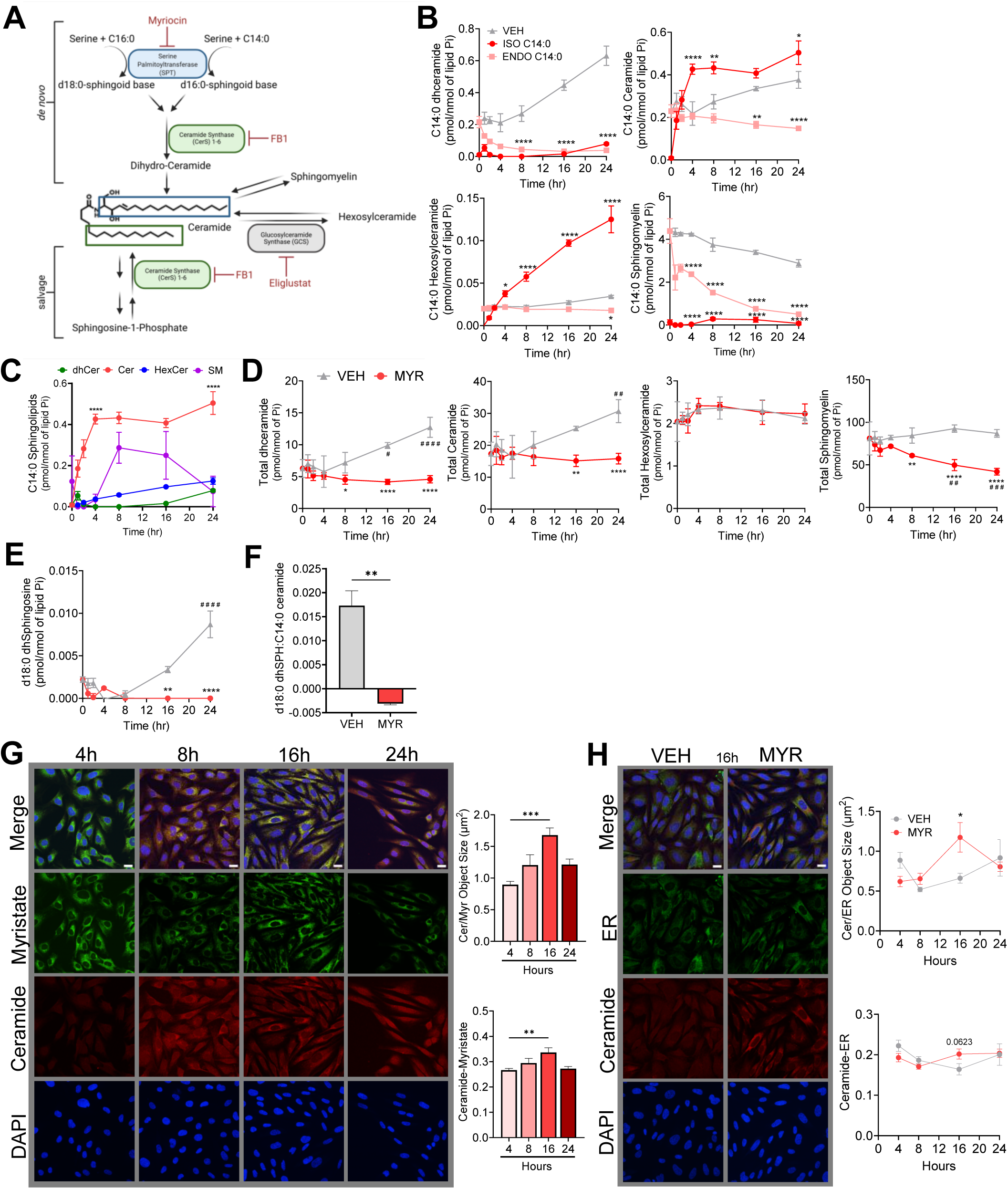
Myristate increases C14:0 ceramide in the ER. **(A)** Simplified sphingolipid metabolism scheme. Metabolites in black, enzymes and related ceramide components share color coding, and inhibitors in red. **(B-F)** HCEC 1CT cells were treated with an isotope labeled myristate (600µM) to measure incorporation into sphingolipids (ISO C14:0) compared to endogenous (ENDO C14:0) and vehicle (VEH, 2% FAF BSA) using LC-MS/MS and normalized to total lipid phosphate (Pi). **(B)** Isotope labeled myristate incorporation in C14:0 dhceramide, ceramide, hexosylceramide, and sphingomyelin compared to VEH over time. **(C)** Isotope labeled myristate (ISO C14:0) in C14:0 sphingolipids over time; ****p<0.0001 for C14:0 ceramide compared to C14:0 dhceramide, hexosylceramide, and sphingomyelin. **(D)** Total dhceramide, ceramide, hexosylceramide, and sphingomyelin. **(E)** d18:0 dhsphingosine. **(F)** Ratio of d18:0 dhsphingosine to C14:0 ceramide. **(G&H)** HCEC 1CT were treated with alkyne-myristate (600µM) for designated times and stained. **(G)** Representative images (left) for myristate (green), ceramide (red), and dapi (blue) and quantification (right) of ceramide and myristate colocalization object size (top) and Pearson’s correlation coefficient (bottom). **(H)** Representative image at 16 h (left) for calnexin (green), ceramide (red), and dapi (blue) and quantification (right) of ceramide and ER colocalization object size (top) and Pearson’s correlation coefficient (bottom); scale bar = 20µm. Data represent mean ± SEM, n=3 (B-F) and n=2 (G&H), *p<0.05, **p<0.01, ***p<0.001, ****p<0.0001 as compared to VEH.. # p<0.05, ## p<0.01, ### p<0.001, #### p<0.0001 as compared to 0 h. Figure 2A was created in BioRender.

### Ceramide synthase inhibition decreases myristate-induced ER stress, but loss of CerS5/6 increases myristate-induced activation of IRE1α

Myristate increased C14:0 ceramide and C14:0 hexosylceramide generation; therefore, we set out to determine if either lipid were required for myristate-induced ER stress in HCEC 1CT. To that end, we used myriocin to inhibit SPT, Fumonisin B1 (FB1) to inhibit all six CerS isoforms, and eliglustat to inhibit glucosylceramide synthase (GCS) (Figure 2A). FB1, but not myriocin or eliglustat, reduced myristate-induced expression of ERDJ4 and IL-6, but not CHOP (Figure 3A). Interestingly, myriocin intensified myristate-induced ERDJ4 expression, while eliglustat intensified myristate-induced CHOP and IL-6 expression (Figure 3A). FB1 diminished myristate-induced expression and phosphorylation of IRE1α, phosphorylation of JNK (a signaling branch downstream of IRE1α), and XBP1s (Figure 3B, 3C). These data suggest that myristate does not induce ER stress via increased *de novo* incorporation of myristate into the sphingoid base; but rather that CerS, via the salvage pathway, are required for myristate-induced ER stress.

**Figure 3:**
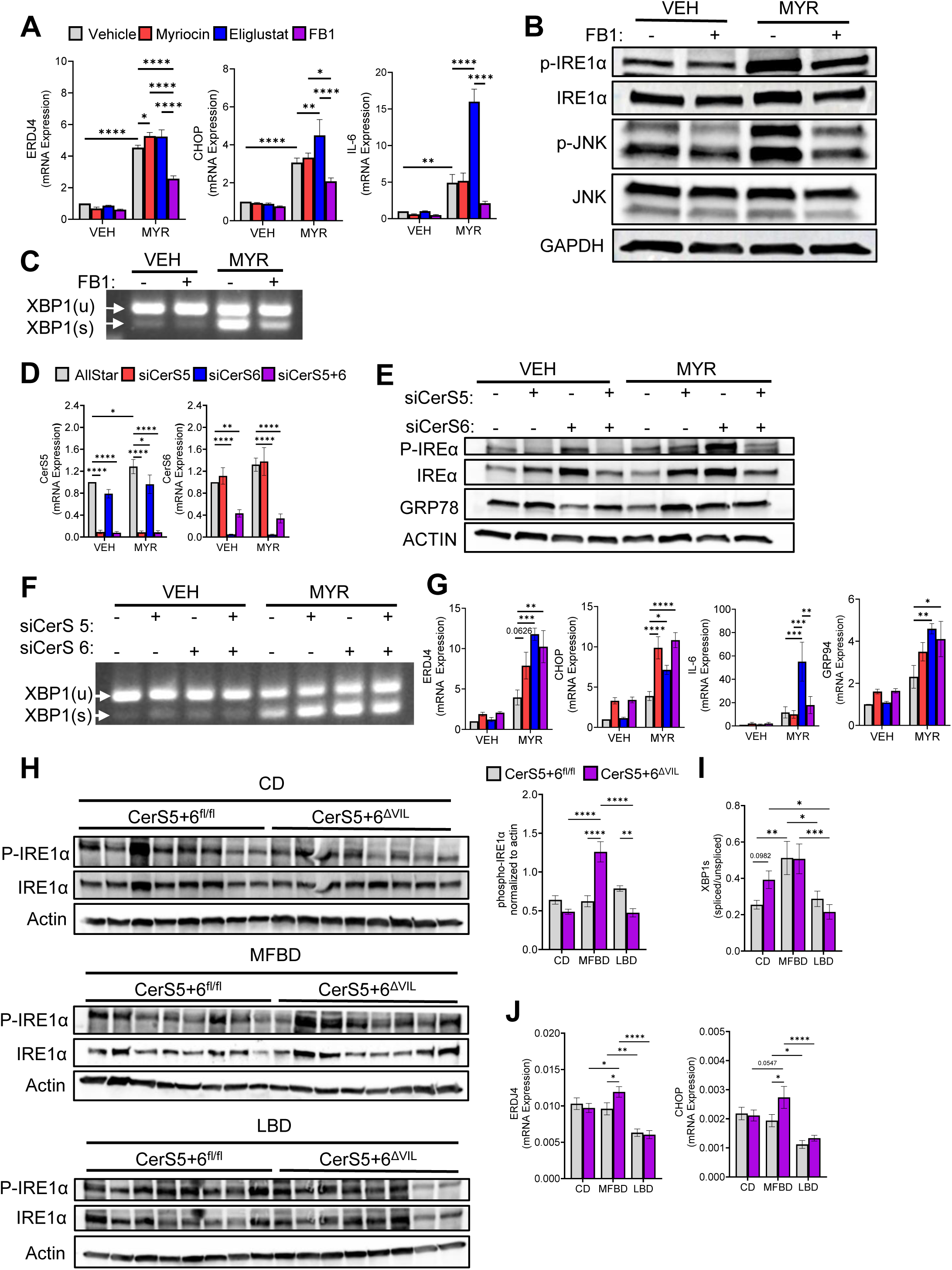
Inhibition of ceramide synthases decreases myristate-induced ER stress, but loss of CerS5/6 increases myristate-induced activation of IRE1α. **(A-C)** HCEC 1CT cells were pretreated with 100nM Myriocin, 100nM Eliglustat, 50µM FB1, or vehicle (methanol) for 1 h then treated with 600µM myristate (MYR) or vehicle (VEH, 2% FAF BSA) for 24 h. **(A)** mRNA expression of ERJD4, CHOP, IL-6 analyzed by RT-qPCR and normalized to RPLPO. **(B)** Western blot of phosphorylated-IREα, total IREα, phosphorylated-JNK, total JNK, and GAPDH as a loading control. **(C)** XBP1 splicing; using extracted RNA and PCR primers to amplify unspliced (XBP1u) and spliced (XBP1s). **(D-G)** HCEC 1CT cells were transfected with 10nM Allstar negative control siRNA or siRNA targeted against CerS5, 6, or both 5 and 6 and treated with 600µM MYR or VEH for 24 h. **(D)** mRNA expression of CerS5 and CerS6. **(E)** Western blot of Western blot of phosphorylated-IREα, total IREα, grp78, and β-actin as a loading control. **(F)** XPB1 splicing and **(E)** mRNA expression of ERDJ4, CHOP, IL-6, and GRP94. **(H-J)** Female and male CerS5+6^fl/fl^ and CerS5+6^ΔVIL^ mice were placed on a 42% milk-fat based diet (MFBD), a 42% lard-fat based diet (LBD), or control diet (CD) for 16 weeks. (H) Western blots of colonic expression of phosphorylated-IREα, total IREα, and β-actin as a loading control (left) and quantification (right). **(I)** Quantification of XBP1 splicing. **(J)** Colonic mRNA expression of ERDJ4 and CHOP were analyzed via RT-qPCR and normalized to actin. Data represent mean ± SEM, n=3 (A-G) and n=6-8 **(H-J)**, *p<0.05, **p<0.01, ***p<0.001, ****p<0.0001.

To determine which CerS isoforms were required for myristate-induced ER stress we used siRNA targeted to CerS5 and/or CerS6 (Figure 3D), the primary isoforms that generate C14:0 ceramide. Surprisingly, knockdown of either isoform exacerbated IRE1α and ATF6-activation after myristate treatment (Figure 3E, 3F, and 3G). To extend our findings *in vivo* and examine the effects of myristate-induced ER stress, focusing specifically on IRE1α and the involvement of Cers5+6 on intestinal biology, we generated mice lacking CerS5 and CerS6 in the intestinal epithelium (CerS5+6^ΔVIL^) and fed them either a 42% milkfat-based diet (MFBD), a lard-based diet (LBD), or a low-fat control diet (CD) (Table 1) (Supplemental Figure 2A). Use of two distinct HFDs provided insight into mechanisms attributable to specific fatty acids, while controlling for overt obesity. Mice on a MFBD or LBD for 16 weeks gained more weight than those on a CD independent of genotype (Supplemental Figure 2B). Similarly, colon length was not impacted by diet or genotype after 16 weeks (Supplemental Figure 2C). Splenomegaly was increased in MFBD CerS5+6^ΔVIL^ mice; however, there were no genotype differences in CD or LBD-fed mice (Supplemental Figure 2D). Investigation of ER stress *in vivo* determined a MFBD, but not an LBD, increased phosphorylation of IRE1α, as well as ERDJ4 and CHOP in colons from CerS5+6^ΔVIL^ mice (Figure 3H, 3J). XBP1s was increased in all MFBD-fed mice, while XBP1 splicing was similar in LBD and CD mice (Figure 3I). Interestingly, loss of either CerS5 or CerS6 alone had no effect on colonic ER stress markers (Supplemental Figure 2E and 2F), suggesting compensation and/or a synergistic effect from the loss of both CerS5 and 6. These unique differences suggest that both CerS expression and resulting sphingolipids are important in intestinal biology through maintaining ER homeostasis.

**Table 1:** *In vivo* dietary information.

|  | Control Diet<br>(CD) |  | Milk-Fat Based Diet<br>(MFBD) |  | Lard Based Diet<br>(LBD) |  |
| --- | --- | --- | --- | --- | --- | --- |
| Kcal/g | 3.6 |  | 4.5 |  | 4.5 |  |
| Protein (% kcal from) | 19.1 |  | 15.2 |  | 15.2 |  |
| Carbohydrate (% kcal from) | 67.9 |  | 42.7 |  | 42.7 |  |
| Fat (% kcal from) | 13 |  | 42 |  | 42 |  |
|  | g/kg | % of fat | g/kg | % of fat | g/kg | % of fat |
| C12 – lauric acid | 1.56 | 3 | 6.3 | 3 | 0.0 | 0 |
| C14 – myristic acid | 5.72 | 11 | 23.1 | 11 | 2.1 | 1 |
| C16 – palmitic acid | 15.8 | 29 | 60.9 | 29 | 48.3 | 23 |
| C16:1 – palmitoleic acid | 0.78 | 1.5 | 3.2 | 1.5 | 4.2 | 2 |
| C18 – stearic acid | 6.76 | 13 | 27.3 | 13 | 27.3 | 13 |
| C18:1 – oleic acid | 14 | 27 | 56.7 | 27 | 81.9 | 39 |
| C18:2 – linoleic acid | 2.08 | 4 | 8.4 | 4 | 37.8 | 18 |
| C18:3 – linolenic acid | 0.26 | 0.52 | 1.1 | 0.52 | 2.1 | 1 |

### Inhibition of ceramide synthase increases dhsphingosine following myristate treatment

CerS inhibition protected from myristate-induced ER stress, while CerS5/6 loss exacerbated ER stress. These unexpected outcomes prompted us to ask if differences in the sphingolipid profile could explain our paradoxical data. To this end, we utilized LC-MS/MS lipidomics to elucidate the mechanism by which FB1 protected from myristate-induced activation of IRE1α. FB1, but not myriocin or eliglustat, decreased C14:0 ceramide (Supplemental Figure 3A) and increased d18:0 dhsphingosine following myristate treatment (Figure 4A). Myristate can also be incorporated into the sphingoid backbone via SPT resulting in d16:0 and d16:1 sphingolipids (Figure 2A). Treatment with FB1 also increased d16:0 dhsphingosine nearly 36-fold following myristate treatment (Figure 4A). These changes in sphingolipids led to an increased ratio of dhsphingosine (d16:0 and d18:0) to C14:0 ceramide in FB1 treated cells (Figure 4B). Knockdown of CerS5, but not CerS6, decreased C14:0 ceramide (Supplemental Figure 3B); however, knockdown of neither CerS5 nor CerS6 increased d18:0 dhsphingosine or d16:0 dhsphingosine following myristate treatment (Figure 4C). The combined depletion of the sphingoid bases with minimal changes in C14:0 ceramide accumulation led to no change in the ratio of dhsphingosine:C14:0 ceramide in cells with loss of CerS5 and/or 6 (Figure 4D). CerS2, CerS5, and CerS6 are the most abundant CerS isoforms in HCEC 1CT (Supplemental Figure 3C). It has previously been shown that loss of CerS5 or CerS6 can lead to increased CerS2 activity and generation of very long chain ceramides [13 46]. However, levels of C16:0 ceramide nor C24:0 ceramide, generated by CerS5/6 and CerS2 respectfully, were altered by loss of CerS 5 and/or 6 (Supplemental Figure 3D). Assessment of sphingolipids *in vivo* revealed that loss of CerS5/6 in the colon decreased C14:0 dhceramide, ceramide, hexosylceramide, and sphingomyelin, while only C14:0 SM was significantly elevated in CerS5/6^fl/fl^ mice fed a MFBD (Supplemental Figure 3E). D18:0 dhsphingosine was increased in CerS5+6^ΔVIL^ regardless of diet; however, this only reached significance in LBD fed mice. Though the ratio of d18:0 dhsphingosine to C14:0 ceramide was increased in all CerS5+6^ΔVIL^ mice (Figure 4F), d16:0 dhsphingosine was not detected in colonic tissue for any diet and/or genotype. These data demonstrate that inhibition of CerS uniquely modulates sphingolipid metabolism; decreasing C14:0 ceramide and increasing levels of both d18:0 and d16:0 dhsphingosine and suggests that these lipids may be protective in abating myristate-induced ER stress.

**Figure 4:**
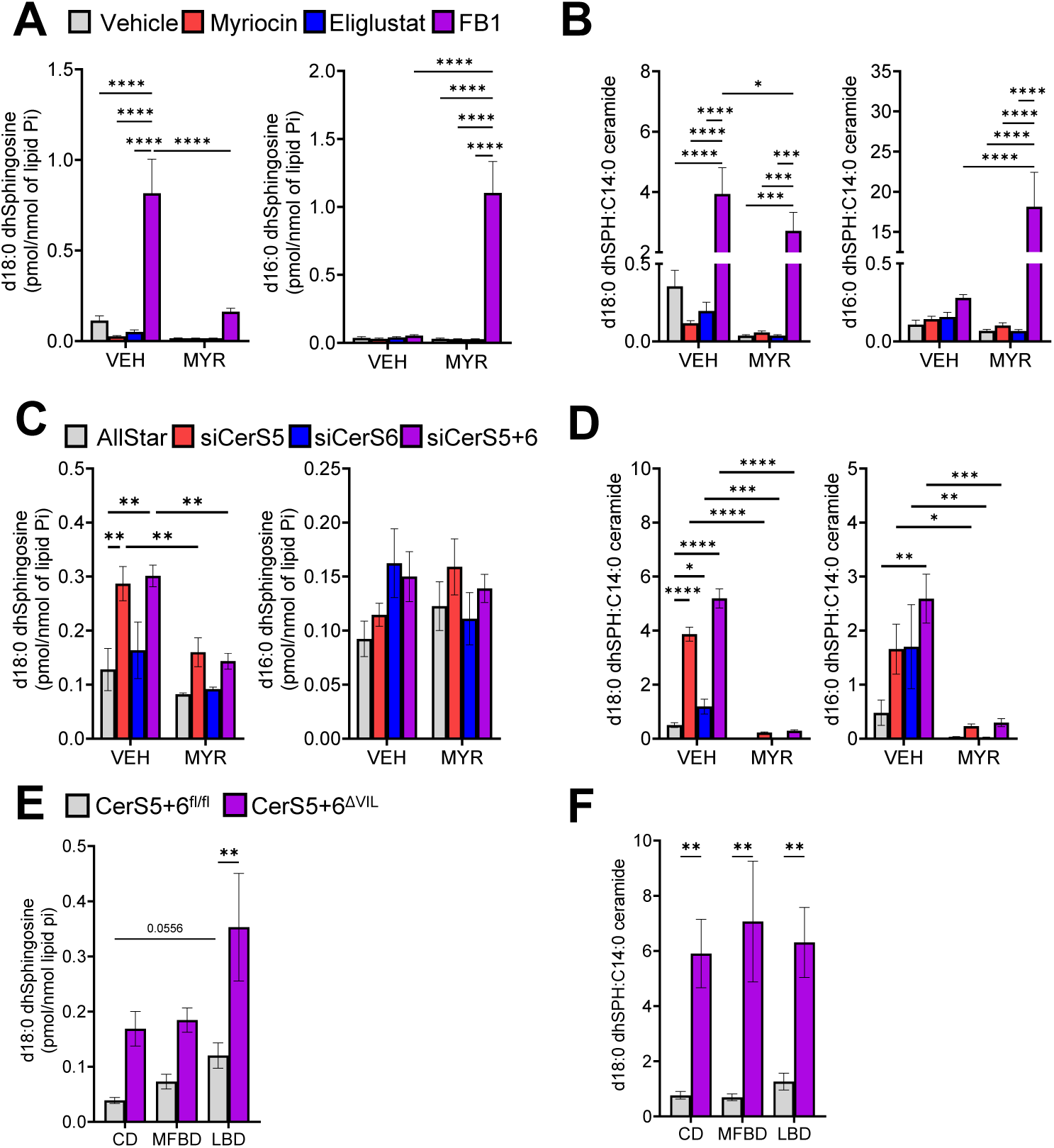
Inhibition of ceramide synthase increases dhsphingosine following myristate treatment. **(A,C,E)** D18:0 and d16:0 dhsphingosine were measured via LC-MS/MS and normalized to total lipid phosphate (Pi). **(B,D,F)** Ratio of sphingoid bases to C14:0 ceramide. **(A&B)** HCEC 1CT cells were pretreated with 100nM Myriocin, 100nM Eliglustat, 50µM FB, or vehicle (methanol) for 1 h then treated with 600µM isotope labeled myristate (MYR) or vehicle (VEH, 2% FAF BSA) for 24 h. **(C&D)** HCEC 1CT cells were transfected with 10nM Allstar negative control siRNA or siRNA targeted against CerS5, 6, or both 5 and 6 and treated with 600µM isotope labeled MYR or VEH for 24 h. **(E&F)** Female and male CerS5+6^fl/fl^ and CerS5+6^ΔVIL^ mice were placed on a MFBD, LBD, or CD for 16 weeks. Data represent mean ± SEM, n=3 (A-D) and n=6-8 (E&F), *p<0.05, **p<0.01, ***p<0.001, ****p<0.0001.

### Accumulation of dhsphingosine protects myristate-induced ER stress

The accumulation of the d18:0 and d16:0 dhsphingosine in cells treated with FB1 led us to interrogate whether these lipids were protective in myristate-induced ER stress. HCEC 1CT were pretreated with myriocin and FB1 in combination to prevent both the accumulation of C14:0 ceramide and the d18:0 and d16:0 dhsphingosine (Figure 2A). As anticipated, FB1 treatment decreased myristate-induced XBP1s and mRNA expression of ER stress and inflammatory markers (Figure 5A, 5B). The addition of myriocin reversed the protective effect of FB1, increasing XBP1s and ERDJ4 expression to similar levels with myristate treatment alone (Figure 5A, 5B). Sphingolipid analyses revealed no differences in d18:0 dhsphingosine in cells treated with FB1 compared to myriocin+FB1 after myristate treatment (Figure 5C). However, d16:0 dhsphingosine was decreased nearly 50% in myristate treated cells preincubated with myriocin+FB1 compared to FB1 alone (Figure 5C). Treatment with myriocin+FB1 decreased the d16:0 dhsphingosine:C14:0 ceramide ratio compared to FB1 alone, while there was no difference in the d18:0 dhsphingosine:C14:0 ceramide ratio (Figure 5D). These data suggest that the accumulation of dhsphingosine, likely d16:0 dhsphingosine, confers partial protection from myristate-induced ER stress.

**Figure 5:**
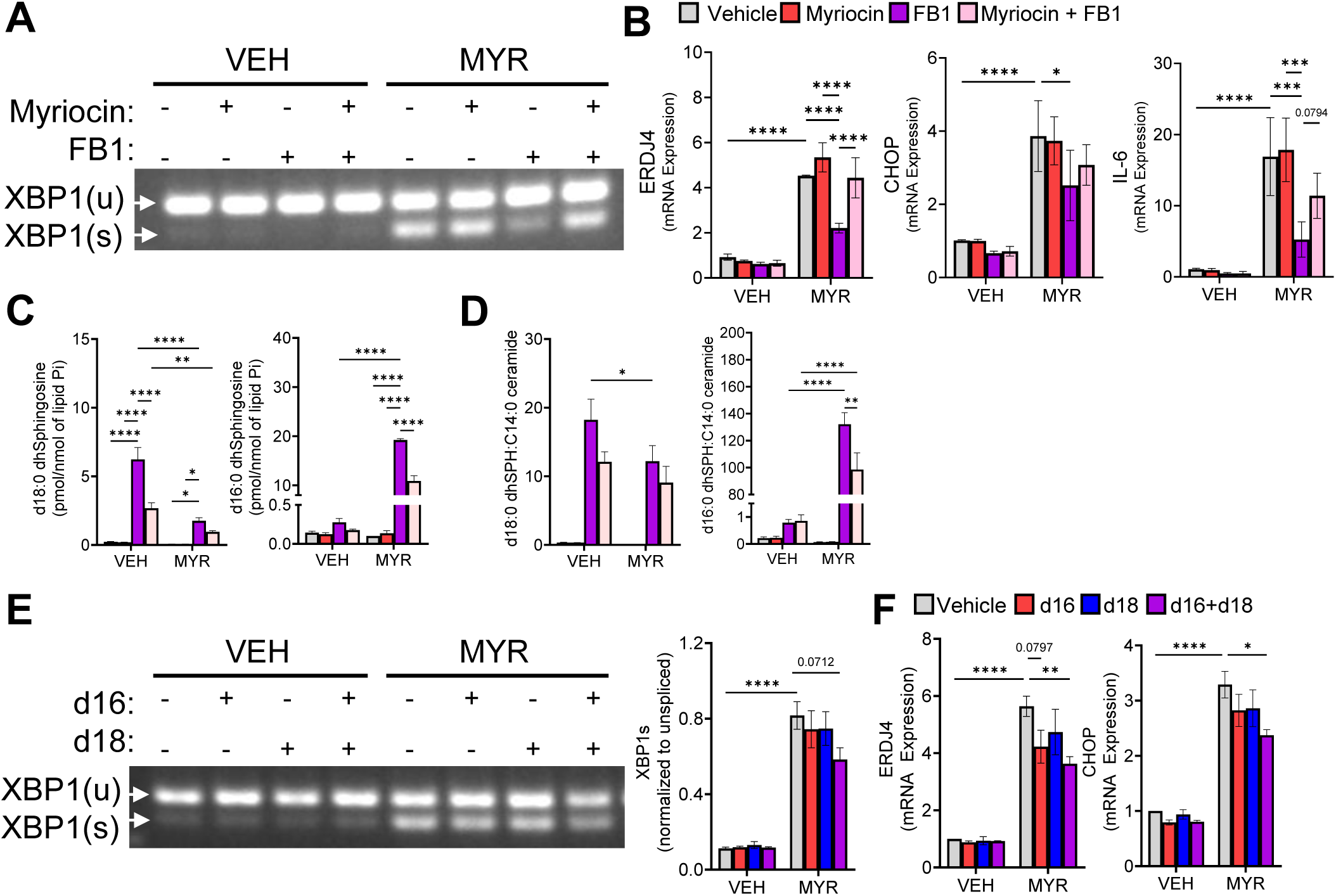
Accumulation of dhsphingosine protects from myristate-induced ER stress. **(A-D)** HCEC 1CT cells were pretreated with 100nM myriocin, 50µM FB1, 100nM myriocin + 50µM FB1, or vehicle (methanol) for 1 h then treated with 600µM myristate (MYR) or vehicle (VEH, 2% FAF BSA) for 24 h. **(A)** XBP1 splicing; using extracted RNA and PCR primers to amplify unspliced (XBP1u) and spliced (XBP1s). **(B)** mRNA expression of ERDJ4, CHOP, and IL-6 were analyzed via RT-qPCR and normalized to RPLPO. **(C)** Sphingoid bases measured via LC-MS/MS and normalized to total lipid phosphate (Pi). **(D)** Ratio of sphingoid bases to C14:0 ceramide. **(E&F)** HCEC 1CT cells were treated with 600µM MYR or VEH for 24 h and 10µM d16:0 dhsphingosine (d16), 1µM d18:0 dhsphingosine (d18), 10µM d16 + 1µM d18 (d16+d18), or vehicle (ethanol) were added to the media for the final 16 hours of incubation. **(D)** XBP1 splicing (left) and quantification (right). **(E)** mRNA expression of ERDJ4 and CHOP. Data represent mean ± SEM, n=3, *p<0.05, **p<0.01, ***p<0.001, ****p<0.0001.

These data led us to probe whether exogenous addition of the sphingoid base could rescue myristate-induced ER stress. To this end, we added d16:0 dhsphingosine, d18:0 dhsphingosine, or both back to the media 8-hours after myristate treatment and assessed activation of ER stress. We measured a trending decrease in XBP1s when both sphingoid bases were added (Figure 5E) and a trending decrease in ERDJ4 expression with the addition of the d16:0 dhsphingosine. The addition of both sphingoid bases generated a small, but significant, decrease in ERDJ4 and CHOP expression (Figure 5F). Together, these data suggest a potential threshold for d16:0 dhsphingosine accumulation, synergistically with d18:0 dhsphingosine, in protecting from myristate-induced ER stress.

## DISCUSSION

HFDs and sphingolipids have consistently been shown to be a major influence on intestinal health and disease [reviewed in [47] and [48]]; however, there has been minimal investigation on the effect of specific fatty acids, enriched in high-fat foods, on sphingolipid metabolism in intestinal pathobiology. Our previous work in rat small intestinal epithelial cells [13] implicated C14:0 ceramide, a ceramide rarely investigated, as a mediator of ER stress and inflammation. In the current study we expanded on these data by examining the role of myristate, a saturated fat found predominately in high-fat dairy, on sphingolipid metabolism and the role of CerS5/6 on ER stress in the colon. Our data revealed that myristate induced accumulation of C14:0 ceramide in the ER along with depletion of dhsphingosine. We hypothesized that the change in the dhsphingosine:C14:0 ceramide ratio induces ER stress. Although inhibition of CerS ameliorated myristate-induced ER stress and inflammation, loss of CerS5 and/or 6 exacerbated IRE1α activation *in vitro* and *in vivo*. Together these data suggest not only species differences in the role of CerS5/6 in the intestines, but that these enzymes may have different functions in the small intestines versus colon.

The difference in CerS inhibition versus knockdown or knockout are perplexing; and contrast with our previous work in which both inhibition and loss of CerS abrogated myristate-induced ER stress. These differences could be due, at least in part, to differences in the models utilized. Our previous work examined myristate-induced ER stress in the context of rat small intestinal epithelial cells, while the current manuscript investigated human colon epithelial cells. Moreover, careful examination as to the differences in sphingolipid metabolism suggests dhsphingosine as a potential mitigator of ER stress in our system. In the time course studies, myristate treatment increased C14:0 ceramide while depleting dhsphingosine; decreasing the dhsphingosine:C14:0 ceramide ratio. Both the increase in C14:0 ceramide and the depletion of dhsphingosine were prevented with FB1. However, knockdown of CerS5, not CerS6, reduced C14:0 ceramide generation with myristate, but d18:0 and d16:0 dhsphingosine were depleted in all knockdown conditions. This led us to hypothesize that d16:0 and d18:0 dhsphingosine are a potential mechanism by which FB1 protected cells from myristate-induced ER stress. We provide novel evidence that modulation of sphingoid bases, either through pharmacological intervention or exogenous addition, conferred partial protection from myristate-induced colonic ER stress. It is possible and likely that generation of C14:0 ceramide and depletion of dhsphingosine both contribute to myristate-induced ER stress. Though canonically due to accumulation of misfolded proteins, ER stress can also be activated by lipid-bilayer stress [37 38]. Investigation into how alterations of these lipids may be contributing to lipid bilayer stress are ongoing. Together, this study highlights the unique mechanism by which myristate (i.e. dairy) impacts intestinal lipid metabolism and ER stress signaling. Further, our results demonstrate that ceramide and dhsphingosine, alongside CerS, are crucial in maintaining intestinal ER homeostasis.

Previous studies in metabolic tissues have demonstrated that ceramides generated from increased fatty acid uptake induced global ER stress [39 41 42 49]. However, these studies focused solely on the fatty acid palmitate. Our current study demonstrates that myristate not only induces ER stress, but specifically activates the IRE1α-XBP1 branch via C14:0 ceramide. Martinez et. al demonstrated that myristate exacerbated palmitate-induced generation of total ceramide accumulation and ER stress in primary mouse hepatocytes [50]. However, in contrast to our study, inhibition of SPT via myriocin blunted the palmitate/myristate ER stress in those cells [50]. Myriocin has also been shown to blunt palmitate-induced ER stress in liver and skeletal muscle [41 49]. In our model myriocin did not decrease activation of IRE1α; however, inhibition of CerS via FB1 abrogated myristate-induced ER stress. Together these studies suggest that different fatty acids may have tissue-specific effects on ER stress and that palmitate and myristate may induce ER stress through different perturbations of sphingolipid metabolism.

Loss of CerS5 and/or CerS6 has previously been shown to alleviate not only ceramide-induced ER Stress [20 39 42], but also ceramide-induced insulin resistance and obesity [21 51]. Our exacerbated ER stress response after loss of CerS was surprising; however, we are not the first to report on the phenomenon [46 52-55]. Others have shown that loss of CerS2, 5, and/or 6 increased ER stress, specifically increasing CHOP expression in breast cancer and neuroblastoma cells lines [46 52]. In our model, loss of CerS5 and/or CerS6 increased activation of both IRE1α and ATF6. In head and neck squamous cell carcinomas (HNSCC) loss of CerS6 induced only ATF6 activation [53 54]. ATF6-activation was attributed to alterations in C16:0 ceramide, release of ER calcium stores, and golgi fragmentation [54]. Although we did not measure golgi fragmentation, C16:0 ceramide levels were not changed by loss of CerS5 and/or 6 compared to control suggesting ER stress activation via a different mechanism. Moreover, the studies in HNSCC did not investigate the role of CerS5 in ER stress. Not only are we the first to report that loss of CerS5/6 induces ER stress in non-cancerous cells and tissues, but our data suggest that CerS and ceramides affect different ER stress pathways depending on the isoform, the lipid, and/or tissue.

Similar to our results, recent work from You et al. demonstrated that HEP2G cells treated with myristate increased C14:0 ceramide and hexosylceramide, decreased total sphingomyelin, and halted *de novo* d18:0 sphingolipid generation [56]. They also reported that myristate treatment alone increased d16:0 sphingolipids globally [56]. However, in HCEC 1CT cells d16:0 dhsphingosine was only increased by myristate following CerS inhibition. Although You et al. did not investigate the cellular effects of d16:0 sphingoid bases, others have begun to implicate them in unique signaling cascades. In heart tissue, a diet high in myristate increased C14:0 ceramide and d16:0 sphingolipids [43 44]. In primary cardiomyocytes exogenous addition of d16:0 sphingolipids, but not d18:0, activated pro-apoptotic signaling via PARP cleavage [44]. Studies have also shown effects of treatment with d16:1 and d18:1 S1P, though not with ER stress. The addition of d16:1 and d18:1 S1P increased the pro-oncogenic connective tissue growth factor (CTGF) via S1P receptor 2 in renal cell carcinoma cells, with d16:1 S1P having higher induction of CTGF than d18:1 [57]. Though our data may seem to contradict these studies, as d16:0 dhsphingosine partially protected from ER stress, studies in Alzheimer’s disease have demonstrated that d16:1 S1P may be protective [58]. In U373-MG cells and mouse primary astrocytes treatment with d18:1 S1P increased IL-1ß and IL-6 expression, while the addition of d16:1 S1P abrogated d18:1 S1P-induced IL-6 [58]. Further, Montefusco et al. recently reported that hepatic loss of SPTLC3, which is responsible for generating d16:0 sphingolipids, lowered fasting blood glucose after HFD-feeding due to suppressed gluconeogenesis [59]. Loss of hepatic SPTLC3 depleted d16:1 sphingomyelin at the plasma membrane, impaired glucagon-stimulated adenylate cyclase activity, and downstream cAMP signaling [59]. Although current studies investigating the impact of d16:0 sphingolipids are limited, it is becoming evident they may play a role in physiology and are an avenue for further investigation.

Sporadic evidence over the last decade has suggested that the sphingoid backbone may directly impact ER stress through unique mechanisms. Park et al. demonstrated in multiple cell lines that in response to ER stress cellular S1P, but not dhS1P, interacted with HSP90a and GRP94 to form a signaling complex with IRE1α, TRAF2, and RIP1 leading to NF-κB activation [60]. In yeast, different sphingoid backbones have been shown to differentially impact IRE1α oligomerization [61]. Dhsphingosine was shown to potentiate DTT-induced IRE1α clustering, while phytosphingosine ablated IRE1α clustering [61]. Further, in HEK293 cells, dhsphingosine and dhceramide activated ATF6 independent of proteotoxic stress or calcium perturbations [62]. Activation of ATF6 by either lipid resulted in unique downstream transcription of lipid metabolic genes that did not occur with proteotoxic-induced ATF6 activation [62]. Although these studies did not investigate d16:0 sphingoid bases, they highlight that various sphingolipid backbones harbor different effects on ER stress signaling. Our study adds to the growing collection of work on sphingoid backbones and ER stress in the context of high-fat diets. The addition of myriocin reversed the protective effects of FB1 on myristate-induced ER stress; however, d16:0 dhsphingosine generation was not completely abolished. These data suggest a threshold of d16:0 accumulation may be necessary to protect from myristate-induced ER stress. Our previous work in IEC6 also implied a threshold of C14:0 ceramide accumulation was needed to induce ER stress [13]. Loss of CerS5 or CerS6 decreased C14:0 ceramide by 25%; however, this was enough to reverse myristate-induced ER stress [13]. Further, exogenous addition of d16:0 dhsphingosine only partially ablated myristate-induced ER stress. Snider et al., previously showed that treatment with a noncanonical d17:0 dhsphingosine above 1µM was not retained intracellularly for time points longer than 1 hour [63]. Considering our exogenous dhsphingosine treatments were 16 hours, the small effect on myristate-induced ER stress could be due to incomplete absorption and/or suggest that the protective effect of d16:0 dhsphingosine is specifically due to *de novo* sphingolipid generation in the ER, rather than extracellular import. Ultimately, what remains unresolved is the mechanism by which d16:0 dhsphingosine protects from myristate-induced IRE1α activation. Determining whether the protective effect of d16:0 dhsphingosine is through direct interaction with ER stress proteins or restructuring of the ER lipid bilayer could be a future area of exploration.

Patients with IBD commonly report that milk and dairy worsen symptoms, with high-fat dairy products most frequently listed [64 65]. Investigation into the role of milk-fat on intestinal health has provided varying results depending on the model utilized. In genetic models of IBD using IL10 or MUC2-deficient mice, high milk-fat feeding exacerbated development of colitis [66 67]. Devkota et al. demonstrated that IL10-deficient mice placed on a 37% milk-fat diet for 24 weeks increased taurine-conjugated bile acids, a rich source of sulfur, which increased colonization of *B. wadsworthia* and led to a robust T_H_1 immune response [66]. Additionally, mice fed a 40% milk-fat diet for 5-weeks and challenged with *C. rodentium* developed more severe colitis compared to a diet high in olive oil fat [68]. On the other hand, in models of chemically induced colitis multiple groups have shown that supplementation of a normal diet with milk phospholipids protects from DSS-induced colitis [69–71]. Wang et al. demonstrated that daily supplementation of milk phospholipids for two weeks prior to DSS-challenge maintained intestinal Notch signaling which increased the number and size of goblet cells, mucins, and antimicrobial peptides compared to DSS alone [69]. Interestingly, Garcia et al. reported that milk-polar-lipid supplementation had sex-based differences on DSS-induced colitis dependent on overall dietary fat content [72]. Supplementation of milk-polar-lipids attenuated DSS-induced colitis under high fat feeding conditions, but exacerbated DSS induced colitis under low fat feeding conditions in male, but not female, mice [72]. Though we did not investigate overt colitis, myristate treatment in HCEC 1CT cells increased expression of pro-inflammatory cytokines, in addition to increased ER stress, both of which have been associated with IBD. Our use of two distinct HFDs further highlight the unique impact of different fatty acids on intestinal health. Although both a MFBD and LBD similarly increased weight gain, only the MFBD increased intestinal ER stress in CerS5+6^ΔVIL^ mice. Together, these studies highlight not only the nuances that various components of milk products play in intestinal health, but that length of diet/supplementation exposure, sex, and overall dietary composition also influence disease outcomes. These studies provided some mechanistic insight (gut dysbiosis [66], Notch signaling [69], and IRE1α activation); however, further investigation is still necessary to tease apart the role of milk-fat in intestinal health and disease.

Dietary habits, particularly high fat foods, have been implicated as a major modifiable risk factor in intestinal health. Although we are not the first to investigate the role of dairy; our unique experimental design has allowed us to dissect more deeply the role of a specific dairy fatty acid (myristate) on intestinal biology. Herein, we’ve provided evidence on a role for sphingolipids in response to diet. Our data suggest that CerS are required for myristate-induced intestinal ER stress and that accumulation of dhsphingosine may provide protection. Future studies could be geared at dissecting the differential impacts of the vital nutrients of dairy (vitamin D, calcium, and potassium) from those of dietary fat on intestinal inflammation and IBD.

## METHODS

### Reagents and Materials

DMEM was purchased from Corning (Corning, NY, USA). Fetal Bovine Serum, glutamax, penicillin-streptomycin, PureLink RNA Mini Kit, qPCR primers, Pierce ECL Substrate, and Pierce BCA Protein Assay Kit were purchased from Thermo Fisher Scientific (Waltham, MA, USA). Myristate (M3128) was purchased from Sigma-Aldrich (St. Louis, MO, USA). Isotope labelled myristate (29463), alkyne myristate (13267), fumonisin B1 (62580), myriocin (63150), eliglustat (21487), sphingoid bases, and mass spectrometry standards were purchased from Cayman Chemical (Ann Arbor, MI, USA). 4µ8C (HY-19707) was purchased from MedChemExpress (Monmouth Junction, NJ, USA). Anti-phospho-EIF2a (9721S), anti-EIF2a (9722S), anti-IRE1α (3294S), and anti-grp78 (3177S) were purchased from cell signaling technology (Danvers, MA, USA). Anti-phospho-IRE1α (NB100-2323) and anti-beta actin (NBP2-76367) were purchased from Novus Biologicals (Centennial, CO, USA). Anti-ATF6 (24169-1-AP) was purchased from proteintech (Rosemont, IL, USA). XBP1 primers were purchased from Integrated DNA Technologies (Coralville, IA, USA).

### Cell Culture

HCEC 1CT and HCEC 2CT (human colon epithelial cells) were provided by Dr. Curtis Thorne (University of Arizona, Tucson, USA). DMEM (Corning, 10-013-CV) was supplemented with 10% heat-inactivated fetal bovine serum, 1% glutamax, and 1% Penicillin-Streptomycin. Cells were kept in a humidified incubator at 37°C with 5% CO2. Cells were seeded and treated as needed 24 h after plating. Cells were grown to a final confluence of ∼75%.

### Animals and Diets

CerS5^fl/fl^ and CerS6^fl/fl^ animals were generously donated by Dr. Jens Brüning (Max Planck Institute, Germany). We crossed CerS5^fl/fl^ and CerS6^fl/fl^ to generate CerS5+6^fl/fl^. Floxed animals were crossed with villin-Cre promoter mice [B6.Cg-Tg(Vil1-cre)997Gum/J; Jackson Laboratory] to generate CerS5^ΔVIL^, CerS6^ΔVIL^, or CerS5+6^ΔVIL^. Animals were maintained in a temperature-controlled environment with a 12/12 h light/dark cycle. Both female and male littermates were aged to 8 weeks before being randomly placed on either a high milk-fat based diet (42% fat), high lard-fat based diet (42% fat), or a control diet (12% fat) (Table 1). Mice were randomized to each diet and fed for on diet for 16 weeks and monitored weekly. Upon euthanasia colon tissues were formalin fixed or flash frozen. All animal procedures were approved by the University of Arizona Institutional Animal Care and Use Committees.

### Fatty Acid Treatment

Fatty acids were prepared as described by Ross *et al* [73]. In brief, myristate was dissolved in 100% ethanol to a concentration of 100 mM and stored at -20°C. Before myristate treatment cells were serum starved for 8 h. Myristate was aliquoted to designated concentrations, dried under nitrogen, and reconstituted using DMEM supplemented with 2% FA-free, low-endotoxin BSA (Fisher, BP9704-100). Myristate was conjugated to BSA via sonication with incubation at 55°C for 15 min, cooled to 37°C, and media was changed to media containing myristate conjugated to BSA for the indicated times. Pretreatments with pharmacologic inhibitors (FB1 50µM, myriocin 100nM, or eliglustat 100nM) were carried out 1 h before the addition of myristate, 4µ8C (100µM) was added at the time of fatty acid treatment.

### CRISPR Cas9 knockout of IRE1**α**

To knockout IRE1α from HCEC 1CT we used the Lentivirus CRISPR system following manufacturers protocol (Addgene, Watertown, MA, USA). Briefly, guide RNAs targeting IRE1α were used to generate HCEC^ΔIRE1α-A^ (Addgene 76408; ACATCCCGAGACACGGTGGT) and HCEC^ΔIRE1α-B^ (Addgene 76407; CCCTCCCTGGAACAAGACGA). To generate lentivirus, plasmids were co-transfected with one of the guide RNAs, Cas9 (Addgene 52962), psPAX2 (Addgene 12260), and PMD.2G (Addgene 12259) into 293FT cells. Virus containing media was harvested and filtered (0.45µm PVDF membrane) after 48 h. HCEC 1CT cells (100,000) were infected with virus in the presence of polybrene (4 µg/ml). After 24 h cells were selected in puromycin (2 µg/ml) for 5 days. Subsequently, cells were plated and expanded from single cell colonies and maintained in normal growth media. Validation of genetic modification was performed via sequencing, RT-qPCR, and western blot.

### Small Interfering RNA

Four hours after plating, cells were transfected with small interfering RNA (siRNA) using Lipofectamine RNAiMax (13778075, Thermo Fisher Scientific) according to manufacturer’s protocol. Thirty-six hours post-transfection media was changed, and cells were treated as indicated. The following Silencer Select siRNA (Thermo Fisher Scientific) were used in this study: CerS5 (S40553) and CerS6 (s48447). Allstar negative control siRNA (1027281) was purchased from Qiagen (Hildan, Germany).

### Immunoblot Analysis

Cells were washed twice with ice-cold PBS then directly lysed in ice-cold RIPA buffer that contained 1:100 protease inhibitor cocktail, phosphatase inhibitor cocktail, and EDTA (Thermo Fisher Scientific). Equal amounts of protein (18-20 mg) were boiled in NuPAGE LDS sample buffer (Thermo Fisher Scientific) and separated by SDS-PAGE (4–15%, Tris-HCl) using the Bio-Rad Criterion system. Separated proteins were then transferred onto PVDF membrane (Sigma-Aldrich) and blocked with 5% nonfat milk in 1X TBST or 5% BSA in 1X TBST (for phosphorylated proteins) for 1 h at room temperature. Primary antibodies (diluted 1:1000 or 1:5000) were then added to membranes and incubated overnight at 4°C. Membranes were washed 3 times with 1X TBST, then incubated with horseradish peroxidase–conjugated secondary antibodies (diluted 1:10,000) for 1 h at room temperature. Membranes were washed 3 times with 1X TBST, incubated with Pierce ECL Substrate, then processed and scanned on Azure Biosystems 500Q. ImageJ (NIH) was used for immunoblot quantification.

### RNA extraction and real time quantitative RT-qPCR

RNA extraction was performed using the PureLink RNA Mini Kit according to the manufacturer’s protocol. 0.5 µg of RNA was used for cDNA synthesis using the qScript cDNA SuperMix (Avantor) according to the manufacturer’s protocol. Real-time RT-PCR was carried out using the Applied Biosystems QuantStudio 3 Real-Time PCR System (Thermo Fisher Scientific). The following TaqMan probes (Thermo Fisher Scientific) were used: human CHOP (Hs00358796_g1), human ERdj4 (Hs01052402_m1), human IL-6 (Hs00174131_m1), human GRP94 (Hs00427665_g1), mouse ERdj4 (Mm01622956_s1), mouse CHOP (Mm01135937_g1), human RPLPO (Hs00420895_gH VIC PL) or mouse Actin (Mm00607939_s1) were used as the housekeeping genes. Cycle threshold (Ct) values were obtained for each gene of interest. DCt values were calculated, and we calculated the relative gene expression normalized to control samples from DDCt values.

### XBP1 splicing

The RT-PCR product of XBP1 were amplified using the following primers: forward XBP1u 5’-TCC GCA GCA CTC AGA CTA CG -3’, forward XBP1s 5’-CTG AGT CCG CAG CAG GTG -3’, and reverse (both XBP1u and XBP1s) 5’-AGT TGT CCA GAA TGC CCA ACA -3’. The thermal cycling profile consisted of 3 minutes at 95°C, 9 cycles at 94°C for 30 seconds, 65°C for 30 seconds (decreasing 0.5°C each cycle), and 72°C 30 seconds, followed by 25 cycles at 94°C for 30 seconds, 60°C for 30 seconds, and 72°C for 30 seconds, with a final 10 minutes at 72°C. After preparation samples were separated on a 3% agarose gel. Optical density (OD) of bands representing XBP1u (141bp) and XBP1s (120bp) were measured by ImageJ (NIH).

### Mass Spectrometry

Lipids from cell culture and tissue were extracted as previously described [13] with analyses conducted by the UACC Analytical Chemistry Shared Resource Core. Data were normalized to total lipid phosphate (Pi) present in the organic phase [74] of the Bligh and Dyer extraction and detected by phosphomolybdate assay [75].

### Microscopy

An alkyne conjugated myristate and click chemistry were used as previously described [76]. In brief, cells were treated with alkyne myristate as indicated. Cells were rinsed twice (PBS), fixed in 4% paraformaldehyde (15 min), washed three times (3% BSA, PBS), permeabilized (0.05% Triton X-100 - 10 min), and washed three times (3% BSA, PBS). Reaction with Azide-Alexa Fluor 488 to label the alkyne myristate used the Click-iT Cell Reaction Buffer Kit (C10641, ThermoFisher Scientific) following manufacturer’s protocol. After, cells were washed three times (3% BSA, PBS), blocked (10% goat serum in 3% BSA, PBS - 30 min), and incubated with primary antibodies overnight at 4°C. The following primary antibodies were used: anti-ceramide (C8104, Sigma-Aldrich) and anti-calnexin (2679T, Cell Signaling). After three washes (3% BSA, PBS), cells reacted with Alexa-Fluor secondary antibodies (568 and 647, ThermoFisher Sceintific) for 1 h, washed three times (3% BSA, PBS), and mounted in Vectashield (H-1900, Vector Laboratories, Newark, CA, USA). Confocal images were acquired on a Nikon AX R Laser-Scanning confocal microscope using a Nikon 60x Plan Apo 1.42NA objective lens.

Nikon NIS-Elements AR (v6.10) software with General Analysis 3 (GA3) was used for image processing and analysis. To achieve accurate cell segmentation that could be automated, each cell from 12 confocal image files (3 files from each timepoint) was manually segmented using the Binary Layers and Binary Toolbar modules. These 12 annotated image files were used to train a SegmentObjects.ai classifier for 1000 iterations using ceramide and calnexin as the original source channels. This SegmentObjects.ai classifier was incorporated into a GA3 workflow, alongside the following image processing functions used to achieve accurate detection of the myristate, ceramide, and calnexin channels: For myristate and ceramide, a low-pass filter (strength = 2px), gamma correction (gamma = 0.6), and rolling ball background subtraction (Myristic Acid radius = 1.2µm and ceramide radius = 2µm) were applied. For calnexin, a rolling ball background subtraction (radius = 2µm) was applied. The Intensity Threshold function was applied to each of these processed channels to generate binary layers for myristate, ceramide, and calnexin. To perform single-cell analysis, the AND binary operation was used between the myristate, ceramide, and calnexin binaries and the trained SegmentObjects.ai cell body binary layer. To quantify measure ceramide-myristate and ceramide-calnexin colocalized binary objects, the AND binary operation was used between ceramide-myristate and ceramide-calnexin binary layers. The Measure Children function was used to quantify colocalized area and ceramide mean intensity within ceramide-myristate and ceramide-calnexin binary layers.

### Sphingoid Base Treatment

D16:0 dhsphingosine (24376, Cayman Chemical) and d18:0 dhsphingosine (10007945, Cayman Chemical) were dissolved in 100% ethanol to a concentration of 20mM and stored at -20°C. Cells were treated as indicated and 8 h after treatment bases were added to media at designated concentrations.

### Statistical Analysis

Statistical analysis was performed using GraphPad Prism 10 (GraphPad Software, La Jolla, CA, USA). Data are presented as mean ± SEM and analyzed by 2-tailed, unpaired Student’s t test when comparing two groups, one-way ANOVA (with Dunnett’s multiple comparisons test) when comparing two or more groups with one independent variable, or two-way ANOVA (with Dunnett’s multiple comparisons test) when comparing two or more groups with two independent variables. Outliers were identified using the Grubbs’ outlier test with an alpha of 5%. Values of P < 0.05 were considered statistically significant.

## Acknowledgements

Lipidomic analyses (Analytical Chemistry Shared Resource Core) as well as microscopy resources and analyses (Microscopy Shared Resource Core) were provided by the University of Arizona Cancer Center supported by the National Cancer Institute P30 CA023074. The graphical abstract (BioRender.com/w018it9) and sphingolipid metabolism scheme (BioRender.com/ven6s6h) were created in BioRender.

## SUPPLEMENTAL METHODS

### MTT Cell Viability Assay

Cells were seeded at densities of 6,000 cells per well in 96-well cell culture plates and treated with increasing doses of myristate, as previously described, for 24 h. Cells then received 10uL of 10mg/mL 3-(4,5-dimethyl-thiazol-2-yl)-2,5-diphenyltetrazolium bromide (MTT, Sigma, M5655-1G) for a final concentration of approximately 1mg/mL/well. After 1h incubating at 37°C, MTT-containing media was removed and replaced with 100uL 100% DMSO (Fisher, ICN19141891). MTT crystals were resuspended by gentle pipetting and quantified via absorbance at 570nm and normalized to vehicle.

## Data Availability

All experimental data, analytical methods, and study materials are available from the corresponding author upon request.

## Abbreviations

ATF6: activating transcription factor 6
CD: control diet
CerS: ceramide synthase
CHOP: C/EBP homologous protein
ERDJ4: endoplasmic reticulum-localized DnaJ homolog 4
ER: endoplasmic reticulum
FB1: Fumonisin B1
GCS: glucosylceramide synthase
HFD: high-fat diets
IBD: inflammatory bowel disease
IRE1α: inositol-requiring enzyme 1 alpha
JNK: c-Jun n-terminal kinases
LBD: lard-fat based diet
MFBD: milk-fat based diet
PERK: protein kinase R-like endoplasmic reticulum kinase
SPT: serine palmitoyltransferase
XBP1: x-box binding protein 1

