## Supplemental Data for "Ceramide Synthases Regulate Myristate-Induced Intestinal IRE1α Activation"

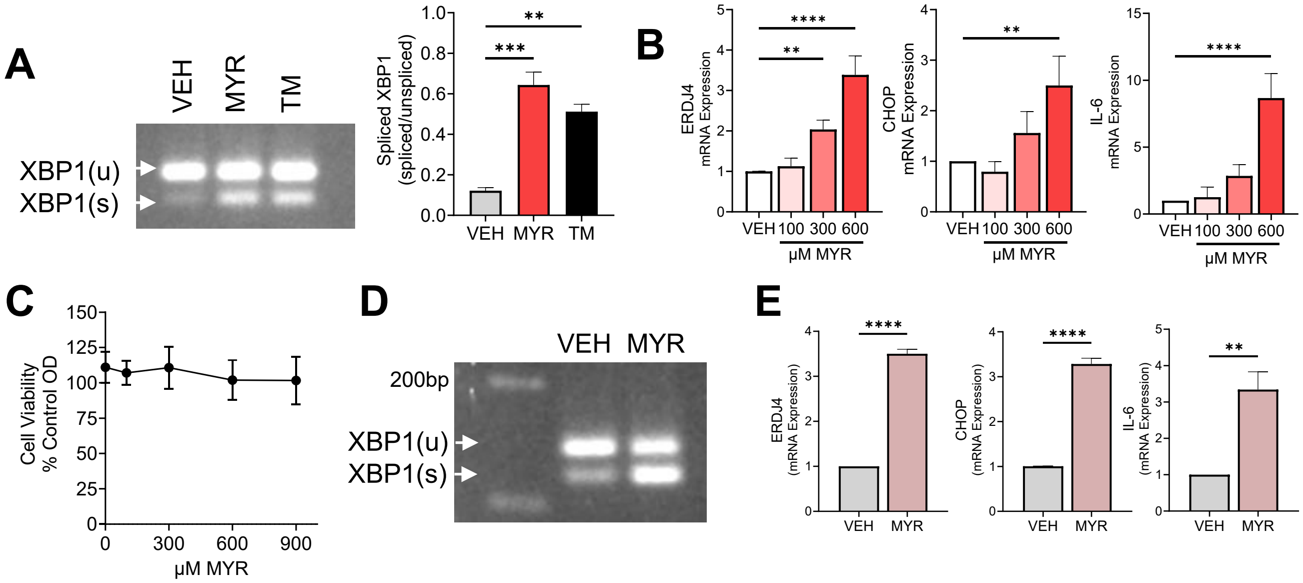


**Supplemental Figure 1: Myristate induces ER stress. (A)** HCEC-1CT cells were treated with 600μM myristate (MYR), tunicamycin (TM), or vehicle (VEH, 2% FAF BSA) for 24 h. XBP1 splicing; using extracted RNA and PCR primers to amplify unspliced (XBP1u) and spliced (XBP1s) with quantification (right). **(B&C)** HCEC-1CT cells were treated with increasing dose of myristate (MYR) or vehicle (VEH, 2% FAF BSA) for 24 h. **(B)** mRNA expression ERJD4, CHOP, and IL-6 analyzed by RT-qPCR and normalized to RPLPO. **(C)** MTT assay to assess cell viability, compared to VEH. **(D&E)** HCEC-2CT were treated with 600μM MYR or VEH (2% FAF BSA) for 24 h. **(D)** XBP1 splicing and **(E)** mRNA expression of ERJD4, CHOP, and IL-6. Data represent mean ± SEM, n=3, $**p<0.01$, $***p<0.001$, $****p<0.0001$.

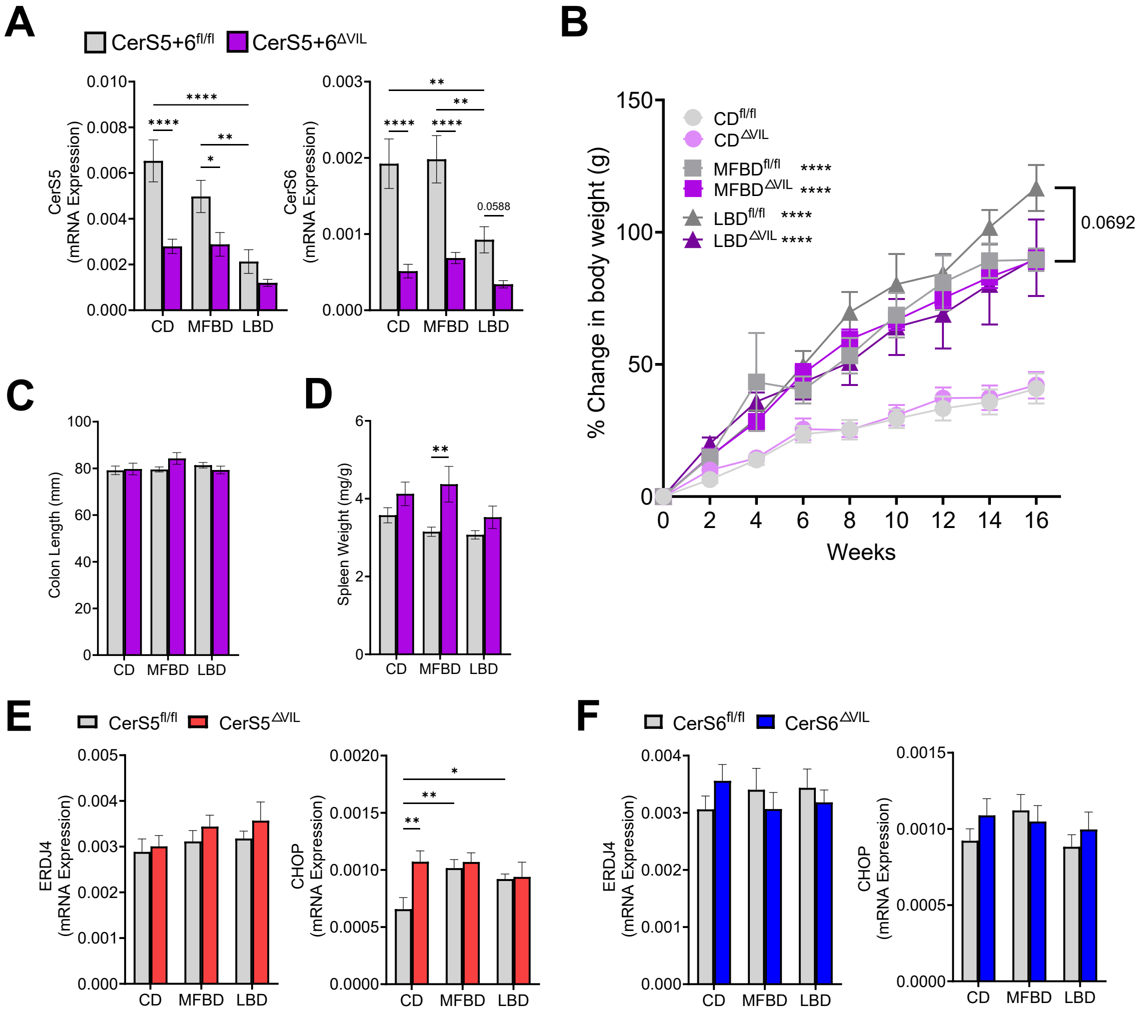


**Supplemental Figure 2: Effects of intestinal loss of CerS5 and/or CerS6.** Female and male CerS5+6$^{fl/fl}$ and CerS5+6$^{\Delta VIL}$ mice were placed on a 42% milk-fat based diet (MFBD), 42% Lard-fat based diet (LBD), or control diet (CD) for 16 weeks. **(A)** Colonic mRNA of CerS5 and CerS6 measured via RT-qPCR and normalized to actin. **(B)** Percent change in body weight of CD, MFBD, and LBD fed CerS5+6$^{fl/fl}$ and CerS5+6$^{\Delta VIL}$ mice. **(C)** Colon length and **(D)** spleen weight normalized to body weight. **(E&F)** Colonic mRNA expression of ERDJ4 and CHOP in **(E)** CerS5$^{fl/fl}$ or CerS5$^{\Delta VIL}$ and **(F)** CerS6$^{fl/fl}$ or CerS6$^{\Delta VIL}$ mice at 16 weeks. Data represent mean ± SEM, n=6-8, *p<0.05, **p<0.01, ***p<0.001, ****p<0.0001.

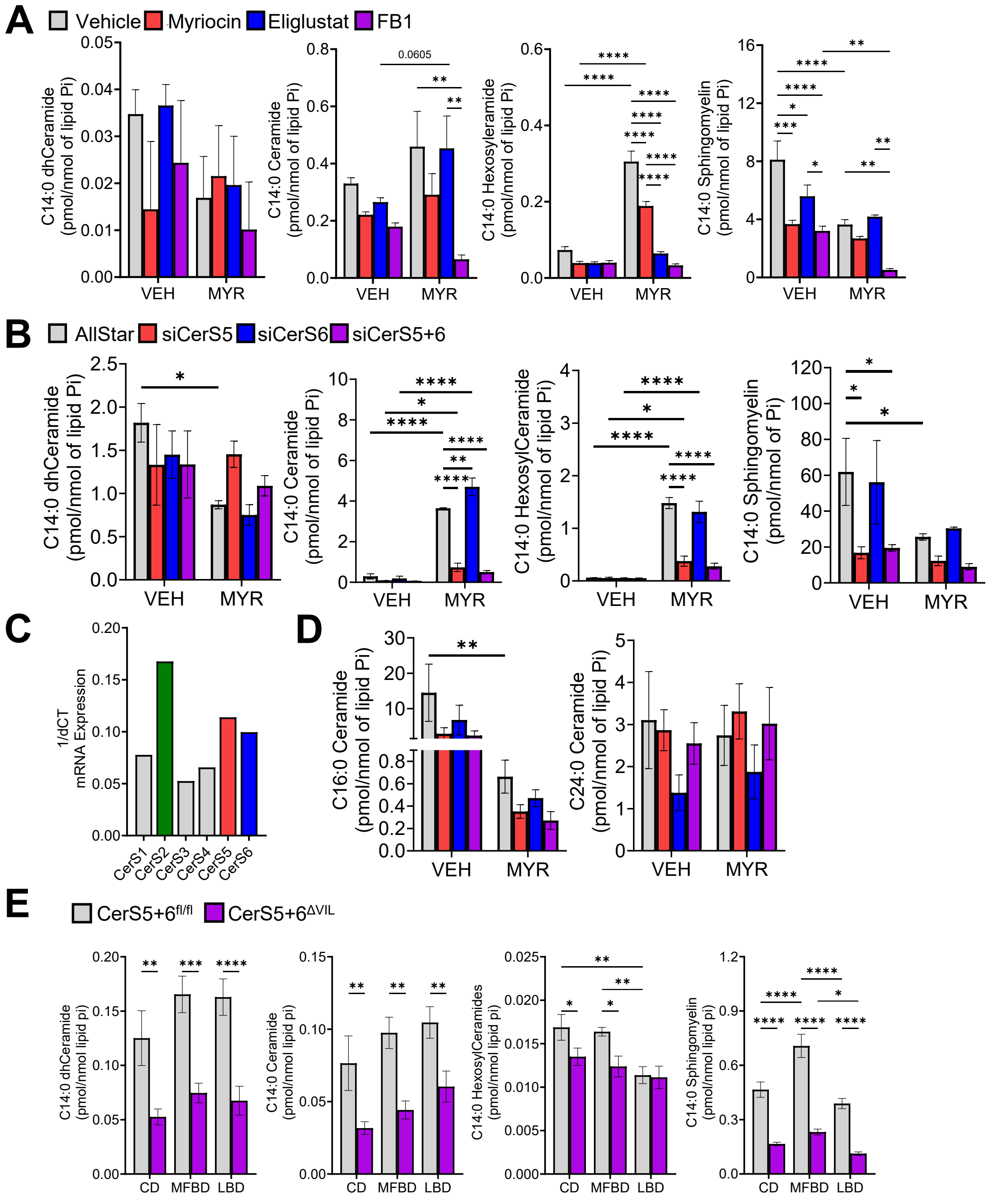


**Supplemental Figure 3: Inhibition versus loss of CerS5/6 uniquely impact C14:0 sphingolipids. (A,B,D,E)** Sphingolipids were measured via LC-MS/MS and normalized to total lipid phosphate (Pi). **(A)** HCEC 1CT cells were pretreated with 100nM Myriocin, 100nM Eliglustat, 50μM FB1, or vehicle (methanol) for 1 h then treated with 600μM of isotope labeled myristate (MYR) or vehicle (VEH, 2% FAF BSA) for 24 h. **(B-D)** HCEC 1CT were transfected with 10nM Allstar negative control siRNA or siRNA targeted against CerS5, 6, or both 5 and 6 and treated with 600μM isotope labeled MYR or VEH (2% FAF BSA) for 24 h. **(C)** mRNA of CerS isoforms measured via RT-qPCR and normalized to actin. **(E)** Female and male CerS5+6$^{fl/fl}$ and CerS5+6$^{\Delta VIL}$ mice were placed on a MFBD, LBD, or CD for 16 weeks. Data represent mean ± SEM, n=3 (A&B) and n=6-8 (D), *p<0.05, **p<0.01, ***p<0.001, ****p<0.0001.
